# Load activated FGFR and beta1 integrins target distinct chondrocyte mechano-response genes

**DOI:** 10.1101/2024.10.11.617817

**Authors:** Helen F. Dietmar, Pia A. Weidmann, Paolo Alberton, Terrilyn Teichwart, Tobias Renkawitz, Andrea Vortkamp, Attila Aszodi, Wiltrud Richter, Solvig Diederichs

**Affiliations:** Experimental Orthopaedics, Research Centre for Molecular and Regenerative Orthopaedics, Department of Orthopaedics, Heidelberg University Hospital, Heidelberg, Germany; Musculoskeletal University Centre Munich (MUM), Department of Orthopaedics and Trauma Surgery, Ludwig-Maximilians-University (LMU), Munich, Germany; Centre for Applied Tissue Engineering and Regenerative Medicine, Munich University of Applied Sciences, Munich, Germany; Division of Hand, Plastic, and Aesthetic Surgery, LMU University Hospital, LMU Munich, Germany; Department of Developmental Biology, Centre for Medical Biotechnology, University of Duisburg-Essen, Essen, Germany; Department of Orthopaedics, Heidelberg University Hospital, Heidelberg, Germany

**Author notes:** Corresponding author PD Dr. Solvig Diederichs Experimental Orthopaedics, Research Centre for Molecular and Regenerative Orthopaedics, Department of Orthopaedics, Heidelberg University Hospital Heidelberg, Germany. Schlierbacher Landstr. 200a 69118 Heidelberg, Germany. These authors contributed equally to this work (Helen F. Dietmar, Pia A. Weidmann).

**Keywords:** Chondrocytes, Mechanotransduction, Fibroblast Growth Factor, Integrins, ERK, Human, Murine, Mouse, Itgb1 knock out

## Abstract

In response to mechanical stimuli, chondrocytes adapt their transcriptional activity, thereby shaping the cellular mechano-response; however, it remains unclear whether the activation of cell surface receptors during mechanical loading converge in the activation of the same mechano-response genes, or whether pathway-specific genes can be defined. We aimed to determine whether load-activated FGF/FGFR signalling and β1 integrin jointly activate ERK and control the same or distinct subsets of mechano-regulated genes. To this end, tissue-engineered neocartilage was generated from murine costal chondrocytes or human articular chondrocytes and subjected to dynamic unconfined compression with or without FGFR inhibition. To assess the role of β1 integrins, neocartilage was generated from embryonic β1 integrin-deficient or wild type costal chondrocytes.

Load-activated FGFR signalling drove ERK activation in murine chondrocytes, and partially also in human chondrocytes, and mechano-response genes could be classified according to their regulation: *Fosl1*, *Itga5, Ngf* and *Timp1* were regulated by load-activated FGFR depending on the developmental stage, whereas β1 integrins controlled *Inhba* expression. In human chondrocytes, load-activated FGFR controlled expression of *BMP2*, *PTGS2* and *DUSP5,* but not *FOSB*.

We show here that the chondrocyte loading response is coordinated by concurrent activation of multiple receptors, and identified for the first time distinct target genes of activated receptors. These insights open up the opportunity to pharmacologically shape the mechano-response of chondrocytes in future studies with promising implications for the management of osteoarthritis and the development of novel therapeutic strategies.

## INTRODUCTION

Cartilage homeostasis, including its biomechanical properties and response to joint loading during normal movement, is disrupted by osteoarthritis (OA), the most common joint disease worldwide ^1–3^. The intracellular signalling cascades activated by mechanical stimulation, the resulting modulation of their target genes and ensuing changes in chondrocyte metabolism remain only incompletely understood in normal as well as diseased cartilage. A detailed understanding of the chondrocyte loading response is important for the development of pharmacological approaches to install and direct a functional loading response in diseased cartilage with mechanical OA aetiologies.

Physical forces are translated into biochemical signals at the cell surface. Bound to heparan sulfate proteoglycans such as perlecan in the cartilage ECM, and released upon mechanical loading, a role as mechano-transducer has been ascribed to Fibroblast growth factor 2 (FGF-2) ^4, 5^. Additionally, integrins have been described as mechanoreceptors in various cell types, including chondrocytes ^1, 6, 7^. In cartilage, mainly collagen-binding integrins (α1β1, α2β1, α10β1), RGD-interacting integrins (αvβ1, αvβ3, αvβ5, α5β1) responsible for fibronectin-binding, and the laminin-binding integrin α6β1 have been described ^8, 9^. Both FGFRs and integrins are known to activate MAP kinases in various contexts ^10, 11^, and the MAPK ERK, p38 and JNK are also activated by mechanical loading of cartilage ^12, 13^. Since ERK has been linked to a variety of important cellular processes like cell proliferation, apoptosis and matrix synthesis, ERK activation is often regarded as a major mechano-transduction event ^4, 12–17^. Previous studies have demonstrated that integrins activate ERK upon mechanical loading of rat chondrocytes ^18, 19^, whereas others have also described that the release of matrix-sequestered FGF-2 by mechanical loading can activate ERK signalling ^4, 5, 20^. Whether load-activated integrin or FGFR are predominantly responsible for ERK activation, or whether their signalling pathways converge on ERK activation remains to be investigated. It also remains unclear, whether models commonly used in the field differ in their mechano-transduction cascades, and whether the same cell surface receptor is responsible for the activation of ERK across those models.

We have previously defined the transcriptomic changes of human articular chondrocytes as well as murine costal chondrocytes that underlie their short-term adaptation to mechanical loading ^21, 22^. The specific mechano-regulated genes differed between human and murine chondrocytes, highlighting the requirement of well-characterised and refined *in vitro* models for cartilage research. Yet, mechano- regulated genes (including *FOS*, *PTGS2*, *DUSP5*, *Ngf*, *Inhba*) that were described to be MAPK/ERK targets ^13, 23–25^, and regulated by FGFR signalling ^26–31^, were identified in both species. Still, it remains unclear whether integrins and FGFRs and their downstream signalling cascades contribute individually to the transcriptomic adjustments that shape the adaptation of cartilage to mechanical load and thus mechano-regulated genes can be categorised into FGFR-dependent and integrin-dependent groups, or whether load-activated integrins and load-activated FGFR converge on ERK activation and jointly control the same mechano-regulated genes. A recent study by Nims et al., deciphered the discrete transcriptomic changes driven by two mechanosensitive ion channels which are activated by distinct loading regimens ^32^; however, it remains to be investigated whether receptor-specific targets or target groups can be identified for other types of receptors, and whether these receptor-specific targets can also be identified within one loading regimen.

We therefore aimed to dissect whether FGFR-driven ERK signalling and integrin-driven ERK signalling control the expression of distinct groups of mechano-regulated genes, in order to determine the relevance of FGFR-driven ERK vs integrin-driven ERK for the transcriptional adaptation to mechanical loading in chondrocytes. To this end, we subjected wild type neocartilage to dynamic unconfined compression in the presence of the FGFR inhibitor PD173074 to determine which mechano-regulated genes are controlled by load-activated FGF/FGFR signalling. Then, we took advantage of an existing, genetically-modified mouse model in which chondrocyte-specific β1 integrin ablation eliminates all major collagen- and fibronectin-binding integrins to address this question ^33^. Following expansion and re-differentiation β1 integrin-deficient and wild type tissue-engineered neocartilage was subjected to a dynamic unconfined compression episode before ERK activation was analysed as well as expression of mechano-regulated genes. Finally, we investigated whether there are FGFR-ERK-dependent and -independent mechano-transductions events also in human articular chondrocytes, by addressing which mechano-regulated genes are controlled by load-activated FGFR-ERK and which are regulated independently of load-activated FGFR signalling.

## MATERIALS AND METHODS

### Transgenic mice and animal husbandry

All experiments were carried out in compliance with the federal and institutional guidelines for the care and use of laboratory animals, approved by the Central Animal Facility of LMU Munich and the government of Upper Bavaria, and approved by the University of Duisburg-Essen and University Hospital Essen. Noon of the day when a vaginal plug was detected was defined as E0.5. *Col2a1*Cre:*Itgb1*^fl/fl^ mice have been described before ^33^ and are labelled in figures as *Itgb1^-/-^*. Animal breedings were approved by the ethics committee of the local authorities (District Government of Upper Bavaria, Germany, Animal Application: TV 55.2-2532.Vet_02-22-161).

### Isolation of murine costal chondrocytes

Primary murine chondrocytes were isolated on E18.5 or P2.5 as described previously ^22, 34^. In brief, single ribs were separated from sternum and spine, digested for 30 min at 37°C in 2 mg/mL collagenase II (Worthington Biochemical, Lakewood, NJ, USA) in DMEM containing 100 U/mL penicillin and 100 µg/mL streptomycin (Gibco^TM^, ThermoFisher Scientific, Darmstadt, Germany) to remove the perichondrium, and finally the remaining cartilage tissue was digested using collagenase solution for 3 h to obtain a single chondrocyte suspension. Cells were expanded in DMEM media containing 10 % FBS (Gibco^TM^, ThermoFisher), 100 U/mL penicillin and 100 µg/mL streptomycin. P2.5 cells were used in P0-P1, E18.5 cells were used in P0. β1 integrin deficient chondrocytes were cultured on vitronectin-coated (R&D, Bio-Techne, Wiesbaden, Germany) culture vessels to facilitate expansion.

### Isolation of human chondrocytes

The study was approved by the local ethics committee on human experimentation of the Medical Faculty of Heidelberg (S-609/2019) and in agreement with the Helsinki Declaration from 1975 (latest version). Articular cartilage samples were obtained from eight donors (five female, two male, 52-88, mean age 71.3 years) undergoing total knee replacement surgery with informed written consent of the patients. Human articular chondrocytes (hAC) were isolated from macroscopically intact cartilage as described previously ^35^. Cells were expanded in DMEM containing 1 g/L glucose, 10 % FBS, 100 U/mL penicillin, 100 µg/mL streptomycin (all Gibco^TM^, ThermoFisher Scientific) for two passages.

### Generation of tissue engineered cartilage

The generation of agarose-based tissue engineered cartilage was described previously ^22^. In brief, cells were collected, counted and resuspended in chondrogenic media at 6 × 10^7 cells/mL. Two volumes of 3 % (w/v) low-melt agarose (Sigma-Aldrich, Darmstadt, Germany) were added, and the suspension cast in custom-made 25 µL moulds to obtain 4 mm x 2 mm discs containing 0.5 million cells. Agarose discs were allowed to solidify at room temperature and cultured in 1.5 mL chondrogenic media (containing DMEM 4.5 g/L glucose, 1 % Penicillin/Streptomycin, 0.1 µM dexamethasone, 0.17 mM ascorbic acid-2- phosphate, 2 mM sodium pyruvate, 0.35 mM proline, 5 mg/mL transferrin, 5 ng/mL sodium selenite, 1.25 mg/mL bovine serum albumin (all Sigma-Aldrich), 5 mg/mL insulin (Lantus, Sanofi-Aventis, Frankfurt, Germany) and 10 ng/mL TGF-β1 (Peprotech, Darmstadt, Germany)). For murine chondrocyte-based neocartilage, ITS+ premix (Corning Life Sciences, Schwerte, Germany) replaced insulin, transferrin and sodium selenite. Medium was changed three times per week.

### Dynamic unconfined compression and FGF receptor inhibition

The dynamic compression protocol was carried out using the software Galil Tools as described previously ^21, 22^. Neocartilage was attached to a glass carrier to mimic the subchondral bone (ROBU, Hattert, Germany) and received fresh media 21 hrs prior to loading. For FGF receptor inhibition, 250 nM PD173074 (abcam, Cambridge, UK) was added 21 hrs before the start of compression and treatment was continued during loading. In brief, engineered tissue was exposed to a single 3 h episode of loading in a custom-designed bioreactor. Initially, the engineered neocartilage was compressed by 10 % of their thickness to ensure the constant contact with the piston during compression. Starting from this 10 % static offset, the engineered neocartilage was compressed by 25 % at 1 Hz during the loading intervals (10 min) whilst maintaining the offset of 10 % during break intervals (10 min). Where indicated, 40 ng/mL FGF-2 (Miltenyi Biotec) was added instead of mechanical loading. After loading, samples were briefly washed in PBS, snap frozen and stored at -80°C. Protein isolation and western blotting Engineered neocartilage was cut in half using a scalpel and minced in 75 µL PhosphoSafe Extraction Reagent (Merck, Darmstadt, Germany), containing 1 mM Pefabloc SC (Sigma-Aldrich) in a mixer mill (Retsch, Haan, Germany) at 30 Hz for 2 x 2 min, with cooling on ice in between cycles. Samples were centrifuged at 13 000 x g for 20 min at 4°C before cell lysates were mixed with 4 x Laemmli buffer and boiled for 5 min at 95°C. Proteins were separated according to size using standard SDS-PAGE and transferred onto a nitrocellulose membrane (Amersham^TM^, GE Healthcare, Chalfont St Giles, UK). Membranes were blocked in 5 % skimmed milk/TBS-T and probed with primary antibodies (listed in supplementary table 1) at 4°C overnight. Detection was carried out using horseradish peroxidase-conjugated secondary antibodies and enhanced chemoluminescence (Roche Diagnostics, Mannheim, Germany/Advansta, USA). Densitometric analysis was performed using ImageJ.

### RNA isolation and quantitative PCR

Engineered neocartilage was halved prior to RNA isolation. Total RNA including miRNA was isolated using a modified QIAquick Gel Extraction Kit and the miRNeasy Mini Kit (Qiagen, Hilden, Germany). In brief, halved samples were minced in 500 µL GQ buffer using a Polytron homogenizer. After incubation at 50°C for 25 min with constant shaking, 750 µL isopropanol was added and RNA was purified using the miRNeasy Mini columns (Qiagen) according to manufacturer’s instructions. For murine chondrocyte-based neocartilage, 166 µL isopropanol was added and RNA was purified using the QIAquick Gel Extraction silica-membrane columns. The Omniscript RT Kit (Qiagen) and oligo(dT) primers were used to transcribe cDNA. Quantitative PCR of cDNA was carried out using SYBR green (ThermoFisher).

Primers are listed in table S2. *Hprt* and *Rpl19* were used as housekeeping genes for murine samples, *CPSF6* and *HNRPH1* were used for human samples and gene expression was calculated using the ΔC_T_ method. Only PCR reactions resulting in a specific band, visualised by agarose gelelectrophoresis, and with the correct melting point were used for data analysis.

### Histology and immunohistochemistry

Halved neocartilage samples were fixed in 4 % paraformaldehyde (PFA) before dehydration in an ascending isopropanol series, and embedded in paraffin wax for cutting. 5 µm sections were cut and de-paraffinized. After rehydration, proteoglycan deposition was analysed by staining 5 µm sections with 0.2 % (w/v) Safranin-O (Fluka, Sigma-Aldrich) in 1 % acetic acid and counterstaining with 0.04 % Certistain Fast Green (Merck) in 0.2 % acetic acid.

Type II collagen immunostaining was carried out as previously established. Briefly, sections were rehydrated, digested with hyaluronidase (Roche Diagnostics, Mannheim, Germany, 4 mg/mL in PBS, pH 5.5), followed by pronase (Roche Diagnostics, 1 mg/mL in PBS, pH 7.4) and blocked in 5 % BSA. After incubation with mouse anti-human type II collagen antibody (see supplementary table 1), sections were incubated with ALP-coupled anti-mouse secondary antibody (see supplementary table 1), and ImmPACT^®^ Vector^®^ Red Substrate (Vector Laboratories, Newark, CA, USA).

### Dimethylmethylene blue assay and Pico Green assay

Halved neocartilage samples were digested in 0.5 mg/mL Proteinase K (ThermoFisher Scientific) in 50 mM Tris-HCl, 1 mM CaCl_2_, pH 8.0 at 65°C with shaking for 18 h. GAG concentration in digests was quantified by addition of the 1,9- dimethylmethylene blue dye using chondroitin sulfate A (Sigma-Aldrich, Darmstadt, Germany) as a standard. DNA contents of digests were measured using Quant-iT- PicoGreen (ThermoFisher) and GAG content was normalised to DNA levels.

### Statistical analysis

The number of biological replicates for each experiment is stated in the figure legend. Statistical analysis was performed in SPSS (Version 29.0.0.0, IBM, Armonk, NY, USA), using a Mann-Whitney U test to assess differences between groups. A *P*-value < 0.05 was considered statistically significant. Where multiple groups were compared, only biologically meaningful comparisons were assessed and a post-hoc Bonferroni-Holm correction applied to adjust for multiple testing.

## RESULTS

### Load-activated FGFR drives ERK phosphorylation, but not gene expression in murine embryonic chondrocytes

To address whether load-activated FGFR signalling is involved in the adaptation of mechano-regulated gene expression, we initially employed engineered neocartilage generated from E18.5 murine costal chondrocytes (as described previously by ^22^). After a pre-cultivation period of 14 days under defined chondrogenic conditions in FGF-free medium, engineered neocartilage was subjected to dynamic unconfined compression in the presence or absence of 250 nM FGFR inhibitor PD173074 (PD). The appropriate PD concentration was selected in a dose-response experiment, in which ERK phosphorylation stimulated by 40 ng/mL FGF-2 was suppressed by 24 hours of treatment with 250 nM PD (Fig. S1). Histological analysis revealed the chondrocyte-typical round cell shape and even distribution of glycosaminoglycans and type II collagen at day 14 of culture in both control and PD-treated neocartilage (Fig. 1A). No significant difference was observed in the GAG/DNA content between PD treated samples and control samples (Fig. 1B), indicating that, as expected, treatment with 250 nM PD for 24 hours at the end of the culture period did not affect overall quality of the engineered neocartilage. As expected, ERK phosphorylation was significantly enhanced by dynamic compression relative to the free-swelling control (Fig. 1C), and also after treatment with 40 ng/mL FGF-2 used as a control. In PD-treated neocartilage, levels of phosphorylated ERK in free-swelling samples were reduced compared to DMSO controls, consistent with endogenous FGF signalling activity in chondrocytes as described previously ^36–38^. In the presence of PD, no significant stimulation of ERK phosphorylation by dynamic compression was observed, demonstrating that load-activated FGFR signalling is a major driver of mechanically induced ERK phosphorylation in murine embryonic chondrocytes. In order to determine how FGFR inhibition, and thus diminished ERK activation, would affect mechano-regulated gene expression, qRT-PCR analysis of our previously defined mechano-responsive genes was performed. Compared to the free-swelling control, *Inhba*, *Fosl1, Itga5, Ngf*, and *Timp1* expression was elevated following compression in the DMSO group as expected (Fig. 1D). Interestingly, FGF-2 treatment also stimulated expression of these genes significantly. Importantly, the load-induced upregulation of *Inhba, Fosl1, Itga5*, *Ngf*, and *Timp1* expression was not inhibited by PD, whereas PD treatment successfully inhibited the stimulation by FGF-2. The load-induced stimulation did not differ significantly between DMSO and PD samples. Thus, whilst mechanical loading activated FGF/FGFR signalling to induce ERK phosphorylation, FGFR-ERK appeared dispensable for the stimulation of mechano-regulated genes in embryonic murine chondrocytes, although all selected genes were FGF-2 targets. Thus, our data demonstrate FGFR-ERK activation is not necessarily required for the short-term transcriptional adaptation to mechanical loading in this model system.

**Figure 1.**
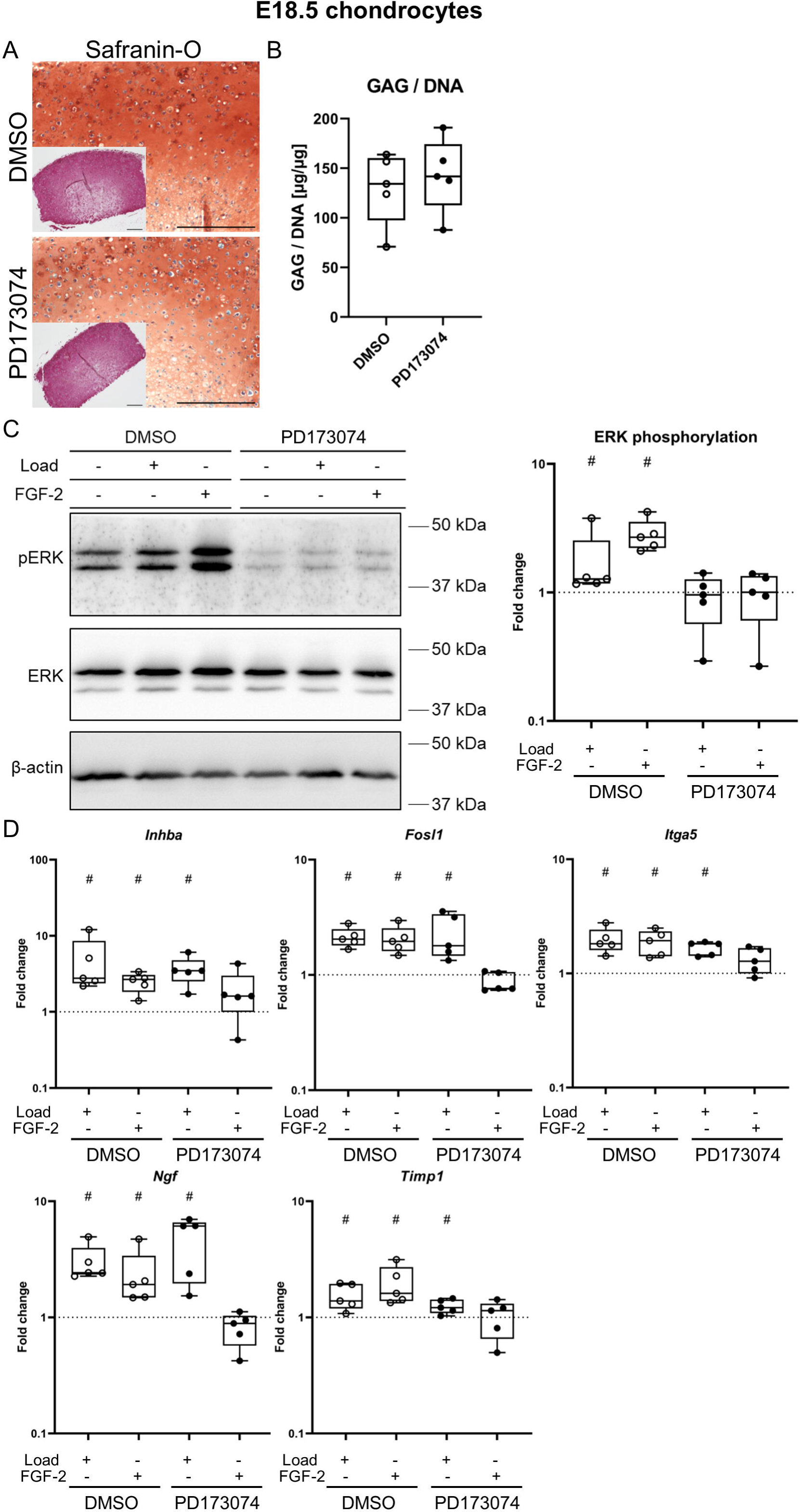
Role of FGFR signalling for mechano-transduction in murine embryonic chondrocytes. Agarose-based neocartilage generated with chondrocytes isolated at E18.5 were cultured for 14 days, then underwent a single, 3 h episode of dynamic unconfined compression in the presence or absence of 250 nM PD173074 (PD) to inhibit FGFR signalling. Additionally, neocartilage was treated with FGF-2 instead of being subjected to compression. (A) Histological analysis of glycosaminoglycan deposition by Safranin-O/Fast Green staining and type II collagen by immunohistochemistry (inset). Scale bar: 500 µm. (B) Glycosaminoglycan content (quantified by DMMB assay) normalised to the DNA content (measured using PicoGreen assay) per neocartilage sample. (C) Visualisation and quantification of ERK phosphorylation by western blotting. Total ERK and β-actin are shown as loading controls. Densitometric analysis of ERK phosphorylation was performed using β-actin as a loading control, and the fold change calculated relative to the free-swelling and non-FGF-2 treated control (dashed line). (D) Gene expression analysis by qRT-PCR, using Rpl19 and Hprt as reference genes. All data are referred to the free-swelling and non-FGF-2 treated controls (dashed line). N = 5, 5 donor populations. Box plots represent the interquartile range (25th – 75th percentile), lines within the box the median, and whiskers extend to the maximum and minimum value. *P*-values were calculated using Mann-Whitney-U test with controls set to 1 and a Bonferroni-Holm correction applied where appropriate. # indicate *p* < 0.05 ctrl vs load/FGF-2, * indicate *p* < 0.05 DMSO vs PD.

### In postnatal chondrocytes load-activated FGFR controls ERK phosphorylation and expression of a subset of mechano-regulated genes

We were surprised by the limited effect of FGFR inhibition on transcriptional regulation since more pronounced effects were reported in other studies using porcine chondrocytes and the same FGFR inhibitor, or an FGF-2 knock-out mouse model ^4, 5, 31, 39^. Thus, we asked whether FGFR signalling would be important for mechano-regulated gene expression after birth. Therefore, we engineered neocartilage as before, this time using murine costal chondrocytes isolated at P2.5. Similar to our observations using E18.5 chondrocytes, treatment with PD did not impair cell morphology or GAG distribution, which however appeared more pericellular than in neocartilage from embryonic chondrocytes (Fig. 2A). Type II collagen deposition and GAG/DNA content of P2.5 chondrocyte neocartilage was also not altered by PD (Fig. 2A, B). Compression as well as FGF-2 treatment still resulted in significantly enhanced phosphorylation of ERK in the DMSO control group, and PD treatment also reduced basal ERK phosphorylation consistent with endogenous FGF activity (Fig. 2C), and allowed no stimulation of ERK during mechanical loading or by FGF-2 treatment. These data demonstrate that load-activated FGFR signalling also drives mechanically induced ERK phosphorylation in P2.5 chondrocytes. When gene expression was examined, relative to the respective free-swelling control, *Inhba, Fosl, Itga5, Ngf,* and *Timp* were upregulated following compression as expected, and also stimulated by FGF-2 treatment similar to our observations in embryonic chondrocytes (Fig. 2D). In agreement with our earlier observations, treatment with PD did not prevent the upregulation of *Inhba* by mechanical loading in 4 out of 5 donor populations. Thus, *Inhba* was again stimulated independently of mechanically activated FGFR signalling. In contrast, the presence of PD during compression prevented a significant upregulation of *Fosl1, Itga5,* and *Timp1.* Whilst *Ngf* expression was still significantly elevated following mechanical loading, this was reduced in the presence of PD. Thus, in early postnatal chondrocytes, load-activated FGFR stimulates ERK phosphorylation, and controls expression of a subset of mechano-regulated genes (*Fosl1, Itga5*, *Timp1* and partially *Ngf*), whereas *Inhba* was not regulated by load-activated FGFR. Therefore, in postnatal chondrocytes, mechano-regulated genes can be classified into FGFR-dependent and FGFR-independent groups, suggesting that modulation of FGFR signalling could allow a targeted manipulation of specific mechano-regulated genes in order to modify the chondrocyte loading response. Whether these FGFR-dependent genes are regulated via ERK remains to be determined using ERK-specific inhibitors, since MEK-independent ERK activation has been described in several cell types, including in response to FGF-2 ^40–43^.

**Figure 2.**
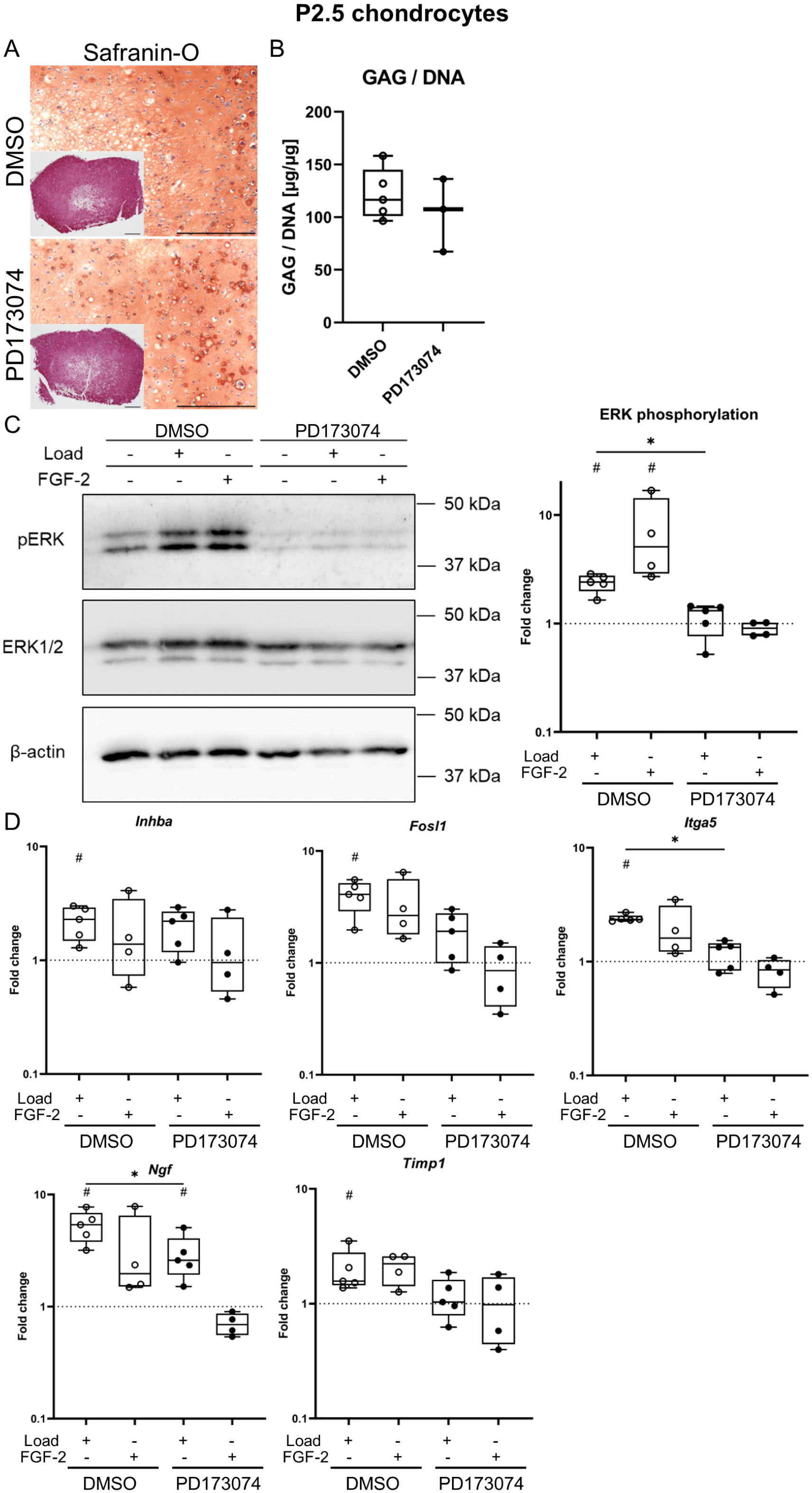
The role of FGFR signalling for mechano-transduction in murine postnatal chondrocytes. Agarose-based neocartilage generated with chondrocytes isolated at P2.5 were cultured for 14 days before undergoing a single, 3 h episode of dynamic unconfined compression in the presence or absence of PD173074 (PD) to inhibit FGFR signalling. Additionally, neocartilage was treated with FGF-2 instead of being subjected to compression. (A) Histological analysis of glycosaminoglycan depositions by Safranin-O/Fast Green staining and type II collagen by immunohistochemistry (inset). Scale bar: 500 µm. (B) Glycosaminoglycan content (measured by DMMB assay) normalised to the DNA content (quantified using PicoGreen assay) per neocartilage sample, n = 5, 4 donor populations (DMSO), 3, 3 donor populations (PD) (C) Visualisation and quantification of ERK phosphorylation by western blotting. Total ERK and β-actin were used as loading controls. Densitometric analysis of ERK phosphorylation was performed using β-actin as a loading control, and the fold change calculated relative to the free-swelling and non-FGF-2 treated controls (dashed line). (D) Gene expression analysis by qRT-PCR, using Rpl19 and Hprt as reference genes. All data are referred to controls (dashed line), n = 5, 4 donor populations (load), n = 4, 4 donor populations (FGF-2). Box plots represent the interquartile range (25th – 75th percentile), lines within the box the median, and whiskers extend to the maximum and minimum value. *P*-values were calculated using Mann-Whitney-U test with controls set to 1 and a Bonferroni-Holm correction applied where appropriate. # indicate *p* < 0.05 ctrl vs load/FGF-2, * indicate *p* < 0.05 DMSO vs PD.

### Without activating ERK, β1 integrins regulate Inhba in murine embryonic chondrocytes

After characterising the transcriptional changes induced by load-activated FGFR, we next assessed the influence of load-activated integrins. Therefore, we employed a chondrocyte-specific deletion of β1 integrin (*Col2a1*:Cre/*Itgb1*^fl/fl^) to genetically eliminate the major integrin types in chondrocytes ^33^. This genetic modification results in a severe perinatal lethal chondrodysplasia associated with reduced chondrocyte proliferation, limiting investigations to embryonic chondrocytes and reducing cell yields from rib cages. In order to obtain enough cells to allow the engineering of neocartilage, we expanded the isolated chondrocytes on the β1 integrin-independent substrate vitronectin and add supporting data from one experiment performed without expansion (data points coloured in red). Unfortunately, the limited number of cells obtained even after expansion also did not permit to include additional experimental groups treated with PD.

We first confirmed the absence of β1 integrin by western blotting, and indeed, β1 integrin was only detected in littermate control, but not *Col2a1*:Cre/*Itgb1*^fl/fl^ chondrocyte-based neocartilage (Fig 3A). Lack of β1 integrins allowed neocartilage maturation as evidenced by GAG deposition according to Safranin-O staining and immunohistochemical detection of type II collagen (Fig. 3B), and quantification of GAG content by DMMB assay revealed no differences in proteoglycan accumulation between wild type and β1 integrin-deficient neocartilage (Fig. 3C). When subjected to dynamic compression, both control and β1-deficient samples exhibited enhanced ERK phosphorylation compared to the free-swelling control (Fig. 3D). This integrin-independent mechanical ERK activation seems consistent with a predominant regulation of ERK phosphorylation by FGFR signalling in embryonic chondrocytes as described above. In order to determine whether mechano-regulated genes are controlled by load-activated integrins, gene expression was analysed in control and β1-deficient neocartilage following dynamic compression (Fig. 3E). After expansion, expression of *Inhba* was elevated in free-swelling β1 integrin-deficient neocartilage, whilst basal expression of *Fosl1*, *Itga5*, *Ngf* and *Timp1* in expanded chondrocytes did not appear different between genotypes (Fig. S2A). Expression of all genes was elevated in free-swelling β1-deficient cartilage neocartilage obtained without expansion. Of note, expression of the selected reference genes was not affected by ablation of β1 integrins (Fig. S2B). Still, the load-induced gene regulation was consistent between expanded and non-expanded chondrocytes, and all tested mechano-regulated genes were induced following compression as expected (Fig. 3E). After mechanical loading, β1 integrin-deficient samples showed no significant upregulation of *Inhba*, despite sustained ERK activation. Expression of *Fosl1* and *Itga5* was stimulated following mechanical loading like in wild type samples. Both *Ngf* and *Timp1* expression was still elevated by mechanical loading in β1 integrin-deficient neocartilage, albeit to a lesser extent than in wild type neocartilage in 4 out of 5 donor populations. Thus, mechano-regulated genes – that were all FGFR-independent in this model – can be grouped into a load-activated β1 integrin-dependent gene (*Inhba*) and integrin-independent genes (*Fosl1* and *Itga5*). *Ngf* and *Timp1* appeared partially, but not mainly, regulated by β1 integrin. Thus, β1 integrins may overlap with load-activated FGFR signalling in the stimulation of *Ngf* and *Timp1*, depending on the developmental timepoint. Importantly, stimulation of *Inhba* expression was controlled by integrins, indicating *Inhba* may be a marker for the activation of β1 integrins by mechanical stimulation. Since stimulation of *Fosl1* and *Itga5* expression was not dependent on integrins or FGFR signalling during mechanical loading of embryonic chondrocytes, their joint activation or a third cell surface receptor must be involved in shaping the transcriptional adaptation of chondrocytes at this developmental stage. Importantly, β1 integrins controlled *Inhba* expression independently of ERK phosphorylation.

**Figure 3.**
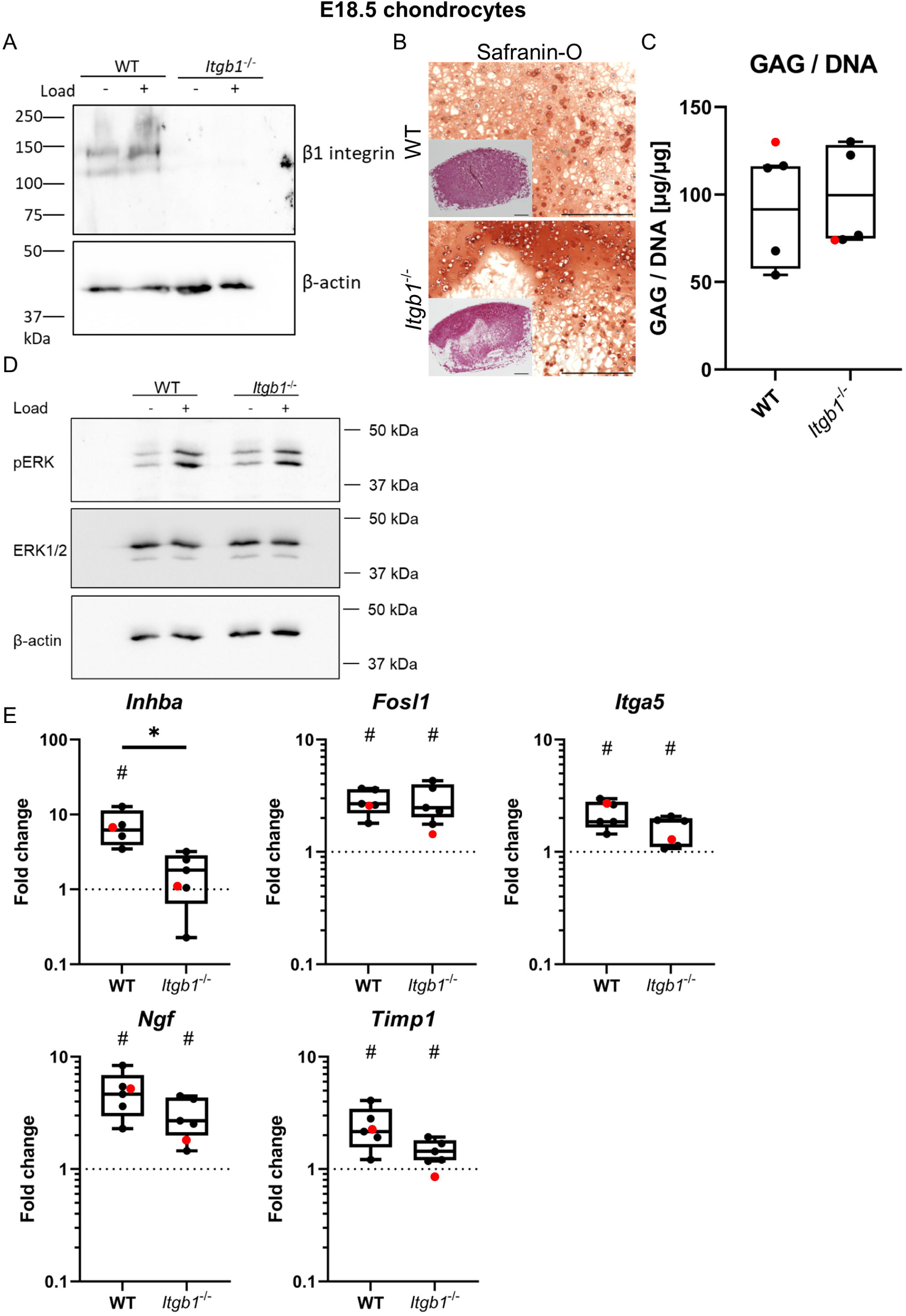
Integrins participate in mechano-transduction in murine embryonic chondrocytes. Agarose-based neocartilage was generated with wild type littermate or *Itgb1*^-/-^ chondrocytes isolated at E18.5, were cultured for 14 days, then underwent a single, 3 h episode of dynamic unconfined compression. (A) Detection of β1 integrin by western blotting. Levels of β-actin are shown as loading control. (B) Histological analysis of glycosaminoglycan deposition by Safranin-O/Fast Green staining and type II collagen by immunohistochemistry (inset). Scale bar: 500 µm. (C) Glycosaminoglycan content (quantified by DMMB assay) normalised to the DNA content (measured using PicoGreen assay) per neocartilage sample. (D) Visualisation of ERK phosphorylation by western blotting. Total ERK and β-actin are shown as loading controls. N = 2 donor populations. (E) Gene expression analysis by qRT-PCR, using Rpl19 and Hprt as reference genes. All data are referred to non- compressed controls (dashed line). Box plots represent the interquartile range (25^th^ – 75^th^ percentile), lines within the box the median, and whiskers extend to the maximum and minimum value. Red data points refer to constructs generated without expansion on vitronectin, n = 5 (expanded, 4 donor populations), 1 (without expansion). *P*-values were calculated using Mann-Whitney-U test with controls set to 1 and a Bonferroni-Holm correction applied where appropriate. # indicate *p* < 0.05 ctrl vs load, * indicate *p* < 0.05 WT vs *Itgb1*^-/-^.

### Load-activated FGFR contributes to ERK activation and drives expression of BMP2 and PTGS2 in human articular chondrocytes

Next, we interrogated in a clinically more relevant model using human articular chondrocytes (hAC) whether load-activated FGFR signalling is also responsible for mechanically activated ERK, whether FGFR-ERK is important for transcriptional adaptation, and whether genes can be classified into FGFR-dependent and - independent subgroups. Therefore, we engineered neocartilage as before, but using hACs obtained after total knee replacement surgery. Histological analysis revealed chondrocyte-typical round cell shape, even glycosaminoglycan distribution and type II collagen deposition throughout the neocartilage after 21 days of maturation, which was not affected by the addition of 250 nM PD for the last 24 hours of culture (Fig. 4A, B). Again, this concentration of PD was effective according to a dose-response experiment (Fig. S3A). Both dynamic compression or treatment with FGF-2 resulted in significantly increased phosphorylation of ERK relative to the free-swelling and untreated control (Fig. 4C). Treatment with PD decreased basal levels of phosphorylated ERK as was observed for murine chondrocyte-based neocartilage, consistent with endogenous FGF activity also in this model system. Remarkably, in hAC-based neocartilage phosphorylation of ERK was slightly but significantly elevated by dynamic compression in the presence of PD, although FGF-2-induced phosphorylation was successfully inhibited by PD. These observations indicate that load-activated FGFR signalling is not the sole driver of mechanically induced ERK phosphorylation in human chondrocytes. In order to determine which mechano- regulated genes are controlled by load-activated FGFR signalling in hACs, and whether PD could suppress their activation despite remaining ERK activation, we analysed expression of *BMP2*, *DUSP5, PTGS2/COX2,* and *FOSB,* which we previously identified as mechano-regulated genes in hACs ^21, 44^. We also examined expression of *ITGA5* and *TIMP1,* since they were mechano-regulated in our murine model (Fig. S3B), but found that they were not regulated by mechanical loading in this cell type, in agreement with our previous work ^21^. As expected, *BMP2*, *DUSP5, PTGS2/COX2,* and *FOSB* expression was upregulated following loading, and interestingly, also stimulated by treatment with FGF-2 (Fig. 4D). Intriguingly, we observed that the stimulation of *BMP2* expression by mechanical loading was strongly reduced by PD. The presence of PD also interfered with the stimulation of *PTGS2/COX2* and *DUSP5* expression following compression. Load-activated FGFR thus drives *BMP2*, *PTGS2* and partially *DUSP5*. *FOSB* expression was below the limit of detection in free-swelling neocartilage in 4 out of 5 donors, but compression and also FGF-2 treatment significantly induced *FOSB* mRNA. Presence of PD did not inhibit load-induced *FOSB* upregulation, but eliminated FGF-2-induced *FOSB* expression. Therefore, also in human articular chondrocytes, mechano-regulated genes could be classified into an FGFR-dependent group (*BMP2*, *PTGS2*, *DUSP5*) and FGFR-independently load-induced *FOSB*. Interestingly, despite incomplete ERK suppression, two out of these three load-activated FGFR-dependent mechano- regulated genes were strongly suppressed, suggesting yet again that ERK may be less important than previously believed for shaping the immediate transcriptional response of chondrocytes to mechanical load.

**Figure 4.**
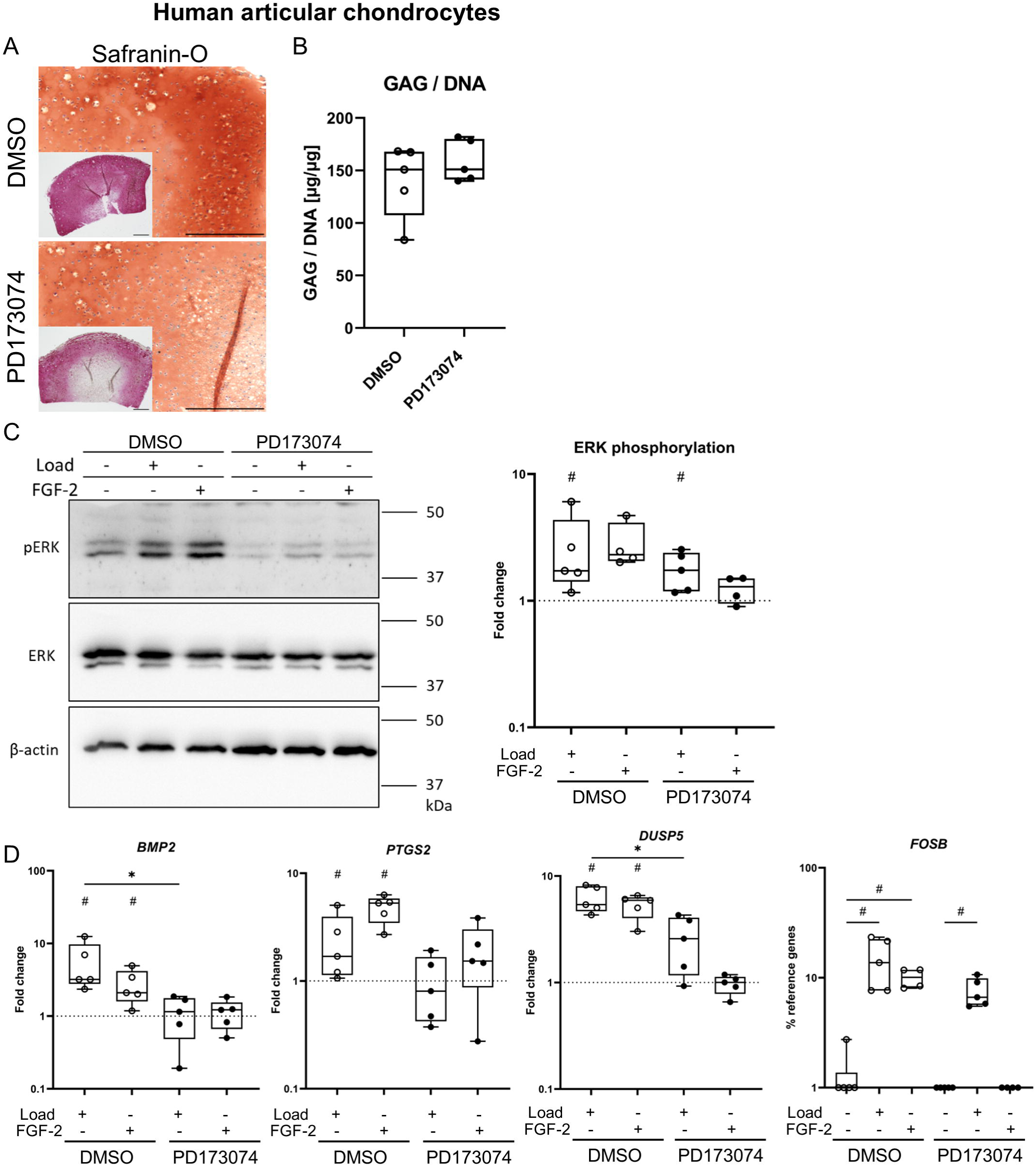
The role of FGFR signalling for mechano-transduction in human articular chondrocytes. Agarose-based neocartilage generated with human articular chondrocytes were cultured for 21 days before undergoing a single, 3 h episode of dynamic unconfined compression in the presence or absence of PD173074 (PD) to inhibit FGFR signalling. In addition, neocartilage was treated with FGF-2 instead of being subjected to compression. (A) Histological analysis of glycosaminoglycans distribution by Safranin-O/Fast Green staining and type II collagen by immunohistochemistry (inset). Scale bar: 500 µm. (B) Glycosaminoglycan content (determined by DMMB assay) normalised to the DNA content (measured using PicoGreen assay) per neocartilage sample. (C) Visualisation and quantification of ERK phosphorylation by western blotting. Total ERK and β-actin were used as loading controls. Densitometric analysis of ERK phosphorylation was performed using β-actin as a loading control, and the fold change calculated relative to the control (dashed line), n = 4-5, 4 donors. (D) Gene expression analysis by qRT-PCR, using *CPSF6* and *HNRPH1* as reference genes. Data are referred to controls (dashed line). For *FOSB*, the relative expression is depicted since expression in control samples was below the limit of detection. N = 5 donors. Box plots represent the interquartile range (25^th^ – 75^th^ percentile), lines within the box the median, and whiskers extend to the maximum and minimum value. *P*-values were calculated using Mann-Whitney-U test with controls set to 1 and a Bonferroni-Holm correction applied where appropriate. # indicate *p* < 0.05 ctrl vs load/FGF-2, * indicate *p* < 0.05 DMSO vs PD.

## DISCUSSION

Appropriate mechanical stimulation is important for articular cartilage maintenance, whilst both lack of loading and overloading can facilitate cartilage degeneration. We have previously defined the short-term transcriptomic response of murine and human chondrocytes to a defined mechanical loading episode ^21, 22^, and herein illuminate for the first time the contribution of load-activated FGFR signalling and β1 integrin signalling, and their induction of ERK to this transcriptional response. Here, we showed that mechano-regulated gene expression is coordinated by the concurrent activation of several cell surface receptors, including load-activated FGFR signalling and β1 integrin, and that they drive discrete events of the chondrocyte loading response. The receptor-specific targets vary in a context-dependent manner, depending on the model system and developmental stages, and we identified here for the first time that FGF/FGFR was responsible for distinct load-activated genes, i.e. *Fosl1*, *Itga5*, and *Timp1* in postnatal murine costal chondrocytes, and in hACs for load-activated *BMP2*, *PTGS2* and *DUSP5*, but not *FOSB*. Technical limitations allowed us to directly compare β1 integrin and load-activated FGFR signalling in only one model system, embryonic murine chondrocytes, in which load-activated FGFR signalling drove mechanically activated ERK, load-activated β1 integrin-controlled load-induced expression of *Inhba* and contributed to stimulation of *Ngf* and *Timp1*, whilst other mechanoreceptors controlled the upregulation of *Fosl1* and *Itga5*. *Inhba* could therefore serve as a marker of β1 integrin-dependent mechano-transduction. Neither load-activated FGFR signalling nor β1 integrins controlled load-induced expression of the remaining mechano-regulated genes, indicating that the concurrent activation of at least one more receptor shapes the short-term loading response of chondrocytes. Taken together, we here demonstrate that load-activated cell surface receptors, including FGFR and β1 integrin signalling do not completely converge in driving the same mechano-transduction pathway and jointly activating the same mechano-response genes, but instead (mechano)receptor-specific response genes can be identified. Whilst long-term effects and a more detailed understanding of receptor-specific events will be addressed in follow-up studies, this new insight opens the possibility to specifically modulate the mechano-response of chondrocytes and pharmacologically activate predominantly anabolic and pro-chondrogenic targets to mimic physiological loading, or inhibit catabolic events in the future.

Whether mechano-response genes classified into β1 integrin-dependent vs FGFR-dependent or -independent groups also exhibit distinct functional aspects for the chondrocyte load response, remains yet to be investigated: The gene products of *Inhba*, Activin A (and Inhibin A) are known to modulate BMP signalling ^45, 46^, an important stimulator of chondrocyte ECM synthesis. Still, conflicting information exist regarding the function of Activin A in cartilage and both, anti-catabolic as well as pro-catabolic effects were suggested ^47, 48^. Thus, it will be highly interesting to address the role of load-induced *Inhba* in cartilage and how the chondrocyte mechano-response can be modified by targeting β1 integrin-*Inhba*.

With *BMP2* and *PTGS2*/*COX2* we identified two genes to be strongly dependent on load-activated FGFR signalling that have already been intensively investigated in chondrocytes. Whilst to our knowledge, the FGFR-mediated mechano-induction of both *BMP2* and *PTGS2* has not yet been reported in human chondrocytes, it is in line with FGF-2 induced *Bmp2* and *Ptgs2* expression during embryonic bone development in mice and in chick osteoblasts, or in murine cartilage and other tissue types, respectively ^29, 31, 49–51^. Also, the stimulation of *PTGS2* following mechanical loading is well described ^21, 44, 52^. *BMP2* is a known stimulator of chondrocyte ECM synthesis and thus contributes to a functional adaptation of cartilage to mechanical load. The role of *PTGS2* for cartilage homeostasis is highly ambiguous, with pro- and anti-inflammatory and anti-hypertrophic functions of its product PGE_2_ being reported ^53–56^. Thus, our data suggest load-induced FGFR signalling may be involved in regulating ECM synthesis and hypertrophic degeneration of chondrocytes. We have previously established mechanical loading regimens producing anabolic and catabolic responses ^57^, and whilst it was beyond the scope of the current study, future studies will need to investigate the contribution of FGFR signalling to these distinct outcomes.

Of note, although not all genes were FGFR target genes during mechanical loading, they all responded to treatment with exogenous FGF-2. This is especially interesting in light of the relative success of sprifermin, a human recombinant FGF-18 protein which is currently being tested in clinical trials for the treatment of OA ^58–60^. Whether FGF-18 also stimulates expression of (FGFR-dependent) mechano-regulated genes, and whether this contributes to beneficial effects for cartilage homeostasis, needs to be addressed in future studies along with the functional impact of receptor-specific targets.

We found indications in two of our model systems that load-activated (FGFR-)ERK is not primarily responsible for shaping the immediate transcriptional response of chondrocytes: Firstly, in embryonic chondrocytes, load-activated FGFR stimulated ERK phosphorylation, but did not control mechano-regulated gene expression; and secondly load-activated FGFR-dependent gene expression was suppressed in human articular chondrocytes although ERK activation was sustained to some degree, which indicates that this important mechano-transducer mediates other cellular responses to mechanical load. These data clearly raise the question by which intracellular pathway(s) load-activated FGFR, β1 integrins and other cell surface receptors shape the transcriptional response. Whether load-activated FGFR signalling involves the activation of ERK to alter gene expression in postnatal murine chondrocytes, and whether load-induced *FOSB* expression is mediated by mechanically activated ERK remains to be determined. Additionally, both FGFRs and β1 integrins are known to signal via several intracellular cascades, including the PI3K family, and its signalling target AKT has been observed to mediate mechanical signals in other systems including osteoblasts and human ankle chondrocytes ^61, 62^.

Still, integrins are also known to signal via Src and FAK, and elevated expression of *INHBA* and *PTGS2* was reported to be partially Src tyrosine kinase-dependent following cartilage injury, which releases FGF-2 from the cartilage matrix similar to mechanical loading ^63^. The Nuclear Factor Kappa B (NFκB) pathway is another intracellular signalling cascade that could potentially be regulated by load-activated β1 integrins and/or FGFR ^64, 65^. β1 integrins or FGFR can also activate MAPK other than ERK ^10, 18, 19, 66–69^ and p38 and JNK were previously implicated in mechano-induced transcriptional regulation in bovine cartilage explants ^13^ and in surgical induction of OA ^70^. Thus, future studies should address the roles of other mechano-transducers, including AKT, Src, NFκB, p38 and JNK, in mediating FGFR- and integrin-driven mechano-transduction.

Surprisingly, and although *Fosl1*, *Itga5, Ngf,* and *Timp1* were regulated by mechanical loading at both timepoints, their expression was driven by distinct cell surface receptors in embryonic and postnatal chondrocytes. Indeed, we were surprised to find that even after a total of three weeks in culture, embryonic and postnatal chondrocytes exhibited differential responses to FGFR inhibition during mechanical loading. It is tempting to speculate that these differences in load-activated FGFR-driven gene expression in murine chondrocytes are due to the fact that P2.5 pups breathe on their own and breast-feed, and that this represents a type of loading conditioning to the cells. Somewhat consistent with our observations, the results of previous studies have suggested distinct effects of FGFR3-activating mutations in the prenatal and postnatal cartilage growth plate ^71–73^. There are several possible reasons for these differences: Altered ECM composition might affect the sequestration and release of endogenous FGF ligands, which may also be expressed at different levels between the timepoints. Differences in chondrocyte maturity, FGF ligand expression and FGFR isoforms may also contribute to distinct effects of FGFR inhibition. Since embryonic and postnatal chondrocytes were derived from the same type of tissue, from an inbreed mouse strain, these models are uniquely suited to address what underlies such signalling dynamics in mechano- transduction.

A major limitation of our study is that due to the severe phenotype, the role of β1 integrins for mechano-transduction in postnatal murine and human chondrocytes could unfortunately not be investigated. Therefore, it cannot be determined with certainty whether in postnatal chondrocytes load-activated integrins still contribute to regulation of *Inhba*, and potentially *Ngf* and *Timp1*. Additionally, due to the technical constraints, we were only able to perform a smaller number of experiments using β1 integrin-deficient chondrocytes and due to the limited number of chondrocytes and neocartilage obtained, FGFR signalling could not be inhibited in the β1 integrin-deficient cells and FGFR-integrin signalling crosstalk could not be assessed. It therefore remains unclear, whether some genes are jointly regulated by FGFR and integrin signalling, or whether they are indeed regulated by other established mechanoreceptors such as PIEZO channels or TRPV4, and downstream calcium signalling ^32, 74, 75^. Unfortunately, the use of alternative approaches, including siRNA, but also *in vitro* application of Cre recombinase and integrin-blocking antibodies, are not compatible with the generation of 3D agarose neocartilage. The dense ECM deposited during the culture period prevents the penetration of the neocartilage, and transient siRNA transfection at the beginning of the culture period did not reduce β1 protein for the duration of culture (data not shown). In addition, the human articular chondrocytes used in this study were obtained after total knee replacement surgery from OA patients, and despite using only macroscopically intact cartilage areas for cell isolation, their origin may of course affect their response to mechanical stimulation. Still, the use of OA chondrocytes is common in the field, mainly because normal non-OA chondrocytes are rarely available. Future studies are needed to compare and identify the drivers and dynamics of mechano-signalling in OA and normal non-OA chondrocytes, which could contribute to therapeutic approaches targeting the chondrocyte mechano-response.

In conclusion, we show here for the first time that the load-driven short-term transcriptional response, an important part of cartilage homeostasis and degenerative diseases like OA, apparently does not necessarily require ERK activation and can be categorised into receptor-specific events. Load-activated cell surface receptors only partially overlap in their respective targets; here, we identified discrete targets of FGF/FGFR and β1 integrin signalling. Load-activated FGFR signalling contributed to the chondrocyte loading response by activating ERK in murine chondrocytes, and regulated a distinct subset of genes (*BMP2*, *PTGS2*, *DUSP5*, and *Itga5*, *Fosl1*, *Timp1*) in human and murine postnatal chondrocytes. Stimulation of *Inhba* was identified as characteristic for load-activated β1 integrins in murine chondrocytes, whereas the remaining mechano-regulated genes where controlled by load-activated FGFR signalling and - perhaps even jointly – by at least one more receptor. Our improved understanding of the chondrocyte loading response stimulates follow-up studies aiming at developing novel pharmacological approaches to install the desired transcriptional adaptation whilst preventing degenerative reactions of chondrocytes in response to physiological load.

## Supporting information

Supplemental table 1 - antibodies

Supplemental table 2 - primer sequences

## Acknowledgements

We would like to thank the clinicians of Heidelberg University Hospital for supplying primary patient material, the patients for their generous tissue donations, and Birgit Frey, Elena Tripel and Carina Binder for excellent technical assistance.

## CRediT authorship statement

**Helen Dietmar:** Investigation, Conceptualisation, Formal analysis, Data curation, Visualisation, Writing – original draft, Writing -review & editing. **Pia Weidmann:** Investigation, Formal analysis, Visualisation, Conceptualisation, Writing -review & editing. **Paolo Alberton:** Methodology, Resources, Writing -review & editing. **Terrilyn Teichwart:** Resources, Writing -review & editing. **Tobias Renkawitz:** Resources, Funding Acquisition, Writing -review & editing. **Andrea Vortkamp:** Resources, Funding Acquisition, Writing -review & editing. **Attila Aszodi:** Conceptualisation, Methodology, Supervision, Funding Acquisition, Resources, Writing -review & editing. **Wiltrud Richter:** Conceptualisation, Project administration, Supervision, Funding Acquisition, Resources, Writing -review & editing. **Solvig Diederichs:** Conceptualisation, Formal analysis, Supervision, Resources, Writing – original draft, Writing – review & editing.

## Data availability

All data are available upon reasonable request to the corresponding author.

This article contains supporting information

