## Supplemental Figure Legends for "Load activated FGFR and beta1 integrins target distinct chondrocyte mechano-response genes"

Figure S1: Dose response of murine rib cage chondrocytes to PD173074. Agarose-based cartilage-like constructs were cultured for 14 days under standard chondrogenic conditions. Constructs were incubated with increasing concentrations of PD173074 (as indicated) for 21 h, before they were treated with 40 ng/mL recombinant FGF-2 for 3 h. ERK phosphorylation in response to FGF-2 was analysed by western blotting, using total ERK and β-actin as loading controls.N = 2.

Figure S2: (A) Expression levels of mechano-regulated genes in non-loaded constructs generated from wild type and β1 integrin-deficient chondrocytes. *Rpl19* and *Hprt* were used as reference genes. Box plots represent the interquartile range (25^th^ – 75^th^ percentile), lines within the box the median, and whiskers extend to the maximum and minimum value. Black dots represent β1 integrin-deficient donor populations that were expanded on vitronectin-coated flasks prior to construct generation, red dots indicate constructs were generated immediately after isolation. (B) C_T_ values for the reference genes *Rpl19* and *Hprt* on d0 (begin of construct maturation, after expansion), and after 14 days of culture under chondrogenic conditions in non-loaded constructs (ctrl) and loaded constructs (load) of wild type and β1 integrin-deficient chondrocytes. Box plots represent the interquartile range (25^th^ – 75^th^ percentile), lines within the box the median, and whiskers extend to the maximum and minimum value. Black dots represent β1 integrin-deficient donor populations that were expanded on vitronectin-coated flasks prior to construct generation, red dots indicate constructs were generated immediately after isolation.

Figure S3: (A) Dose response of human articular chondrocytes to PD173074. Agarose-based cartilage-like constructs were cultured for 21 days under standard chondrogenic conditions. Constructs were incubated with increasing concentrations of PD173074 (as indicated) for 21 h, before they were treated with 40 ng/mL recombinant FGF-2 for 3 h. ERK phosphorylation in response to FGF-2 was analysed by western blotting, using total ERK and β-actin as loading controls. N = 3. (B) Expression of *ITGA5* and *TIMP1* in human articular chondrocyte-derived constructs as measured by qRT-PCR. *CPSF6* and *HNRPH1* were used as reference genes. Data are referred to controls (dashed line). Box plots represent the interquartile range (25^th^ – 75^th^ percentile), lines within the box the median, and whiskers extend to the maximum and minimum value. *P*-values were calculated using Mann-Whitney-U test with controls set to 1, * indicates *p* < 0.05 ctrl vs FGF-2.
