## Supplementary figures and images for "Load activated FGFR and beta1 integrins target distinct chondrocyte mechano-response genes"

### Supplemental Figures 1-3

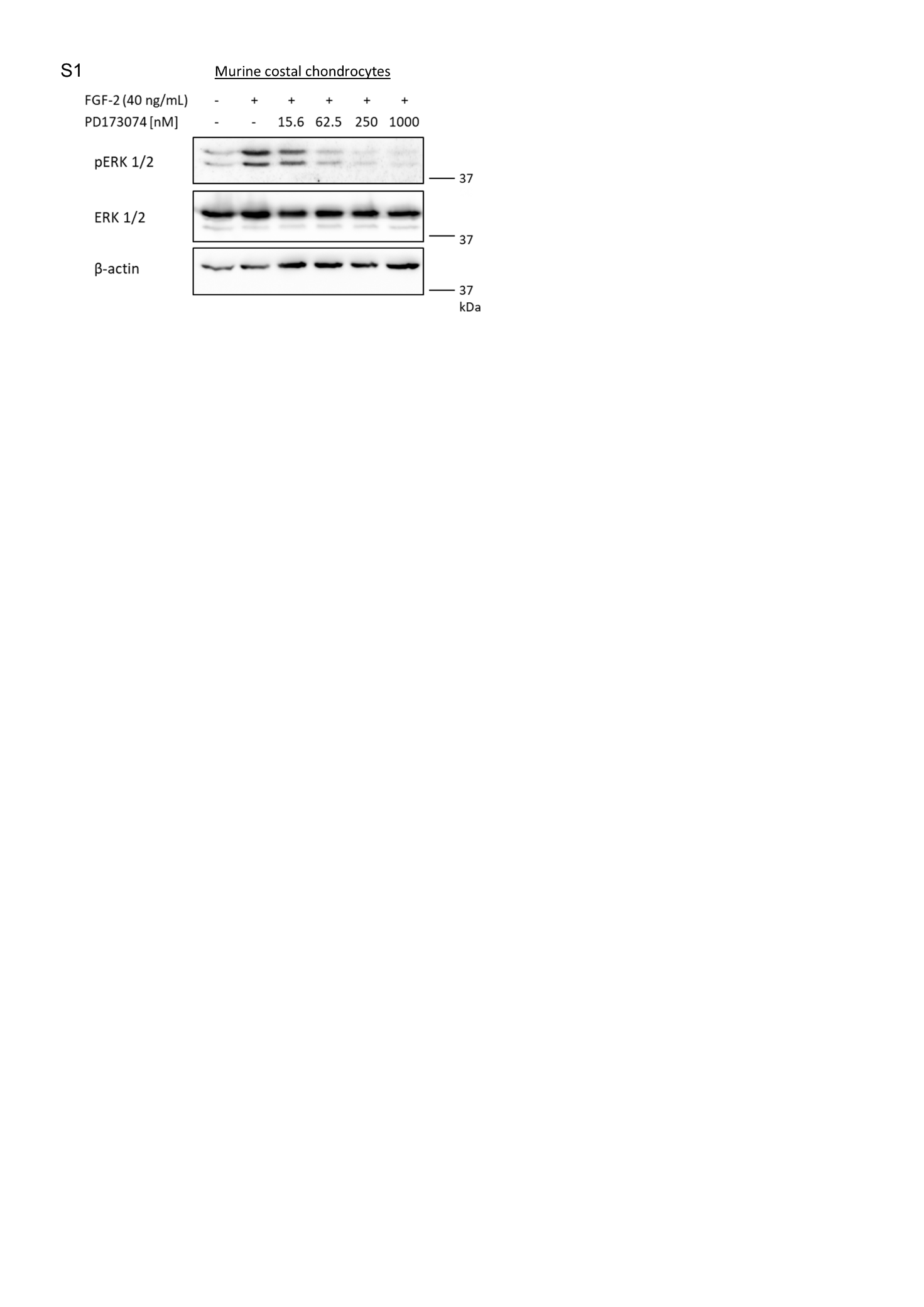

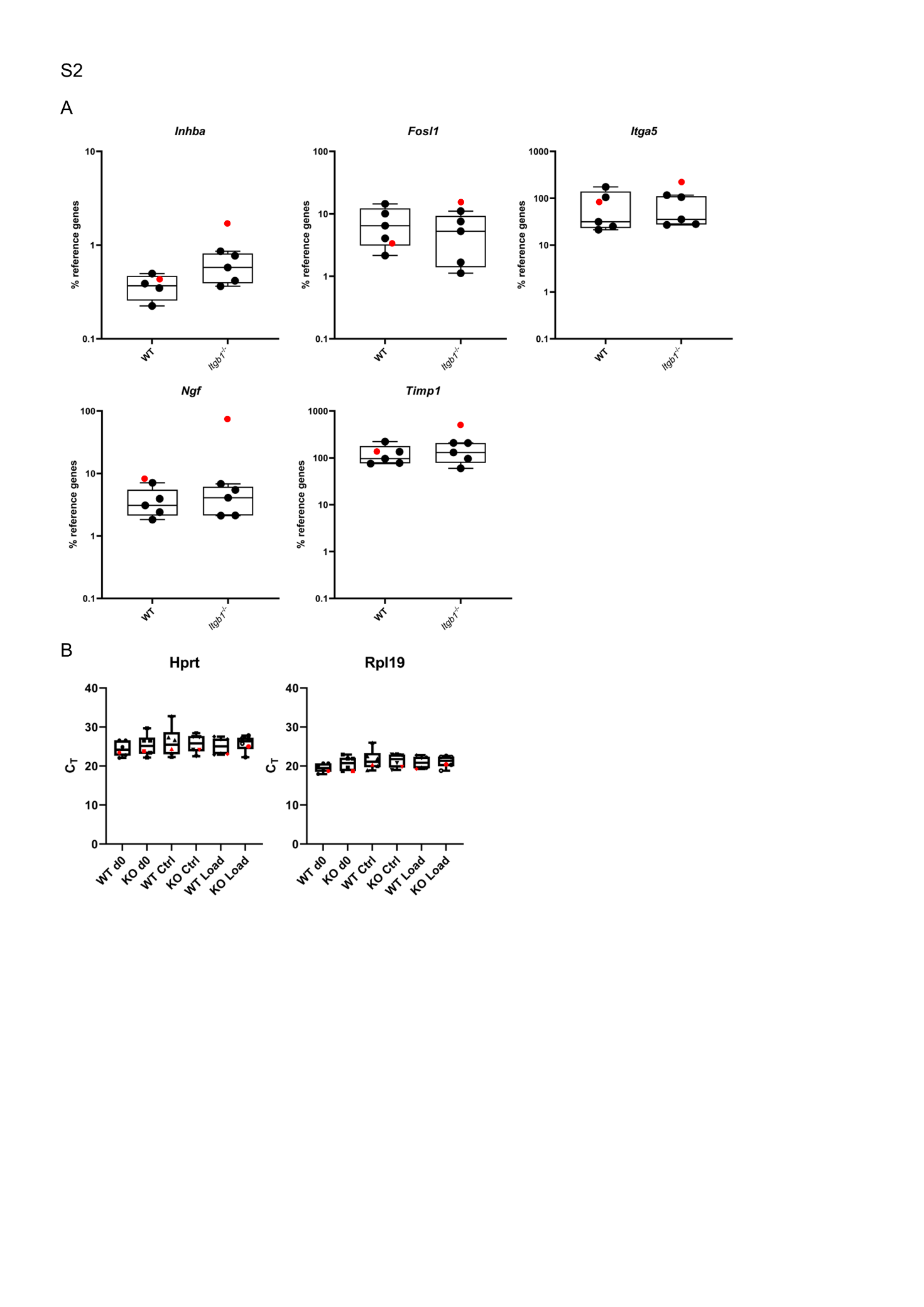

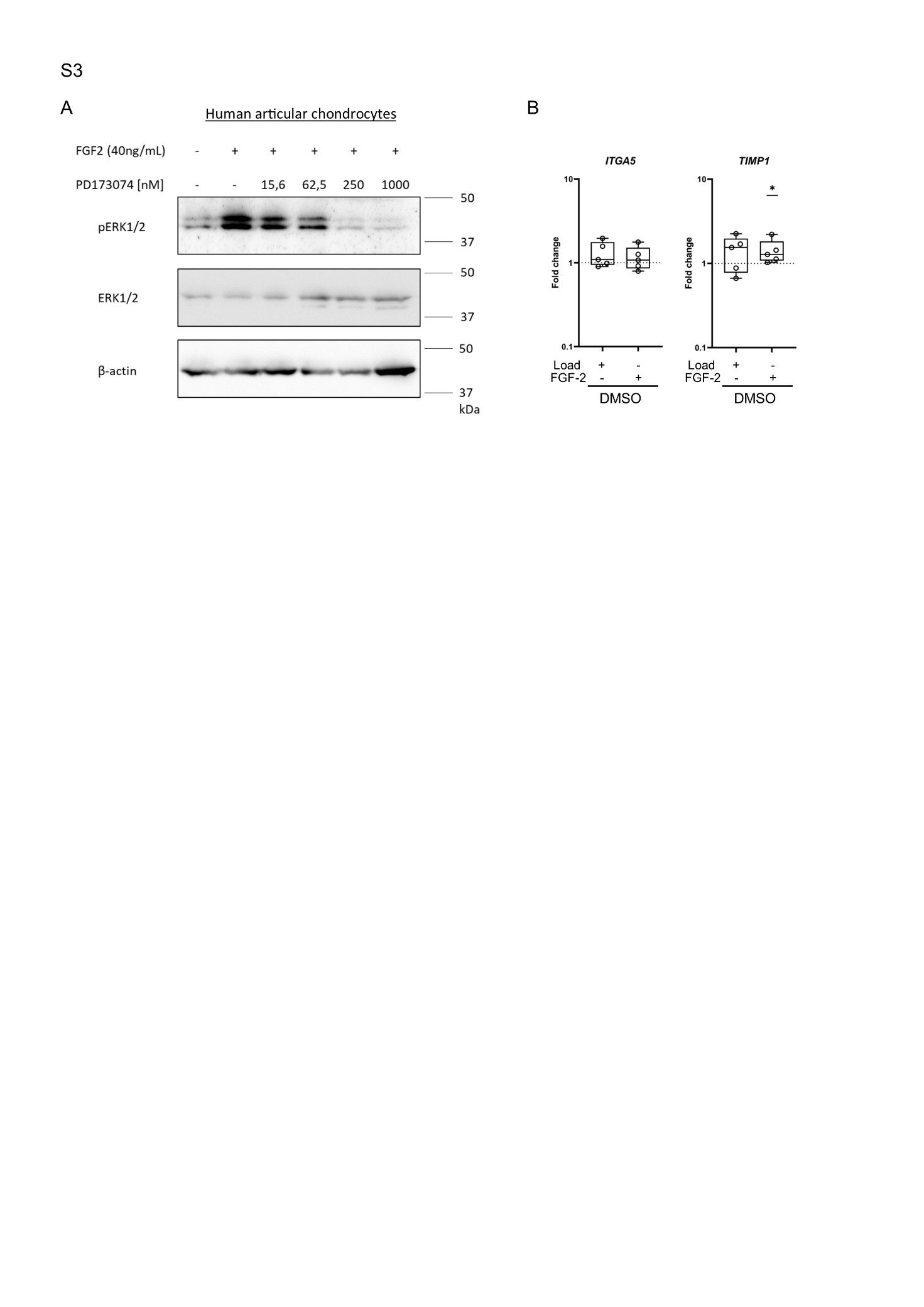

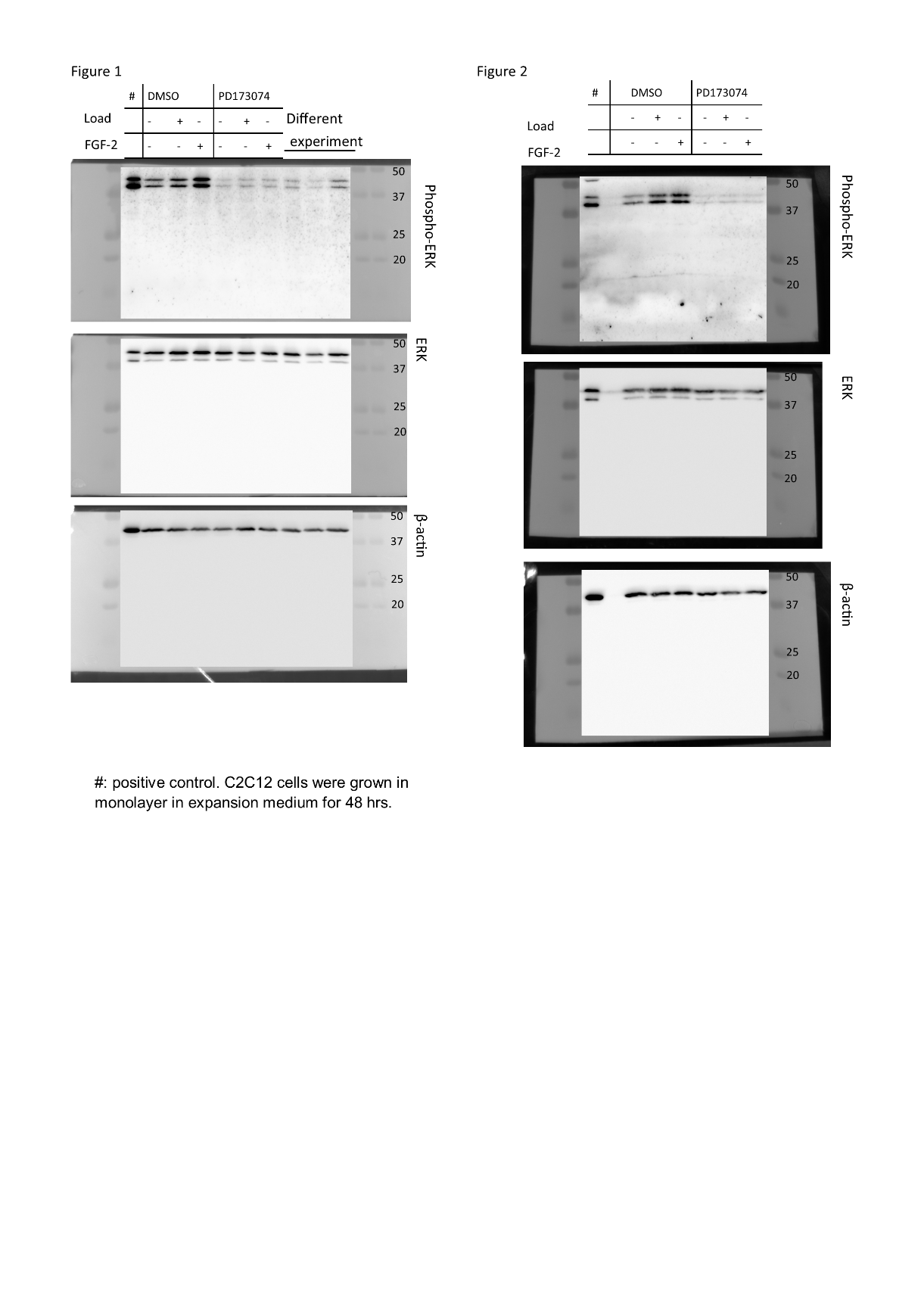

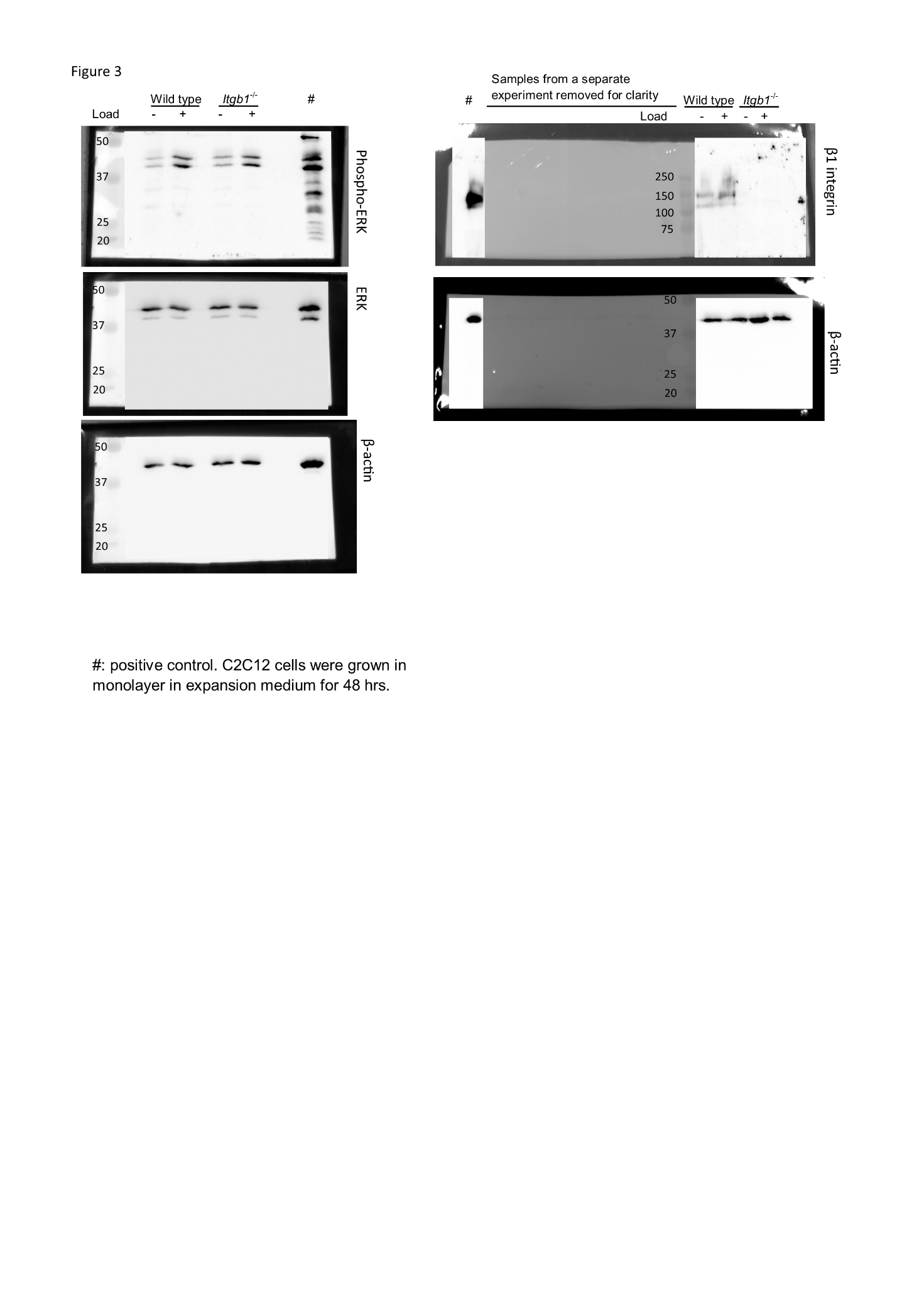

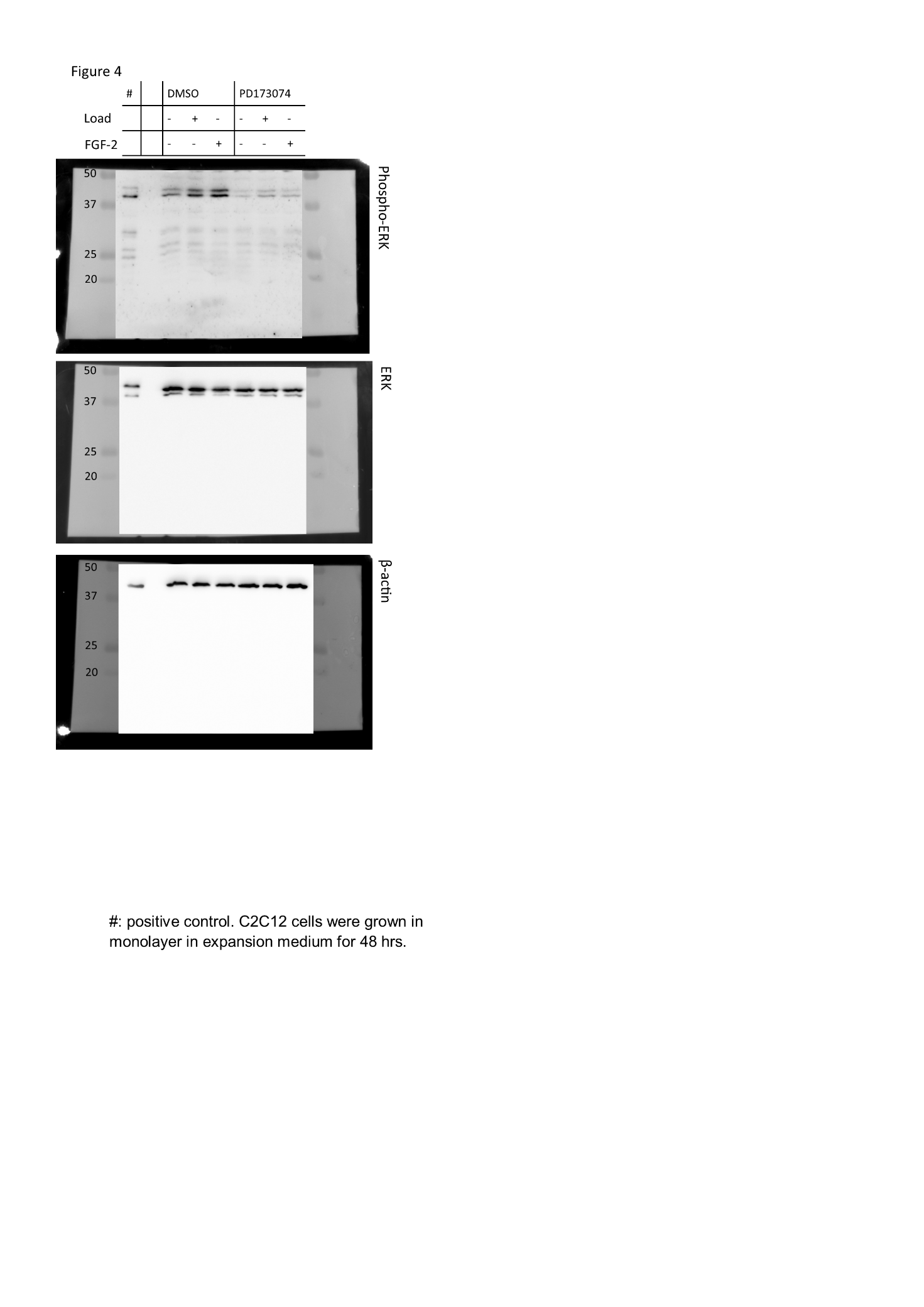
